# The *Synechocystis* sp. PCC 6803 open reading frame *slr0201* that is homologous to *sdhC* from Archaea codes for a [2Fe-2S] protein

**DOI:** 10.1101/2021.09.23.461530

**Authors:** Fusheng Xiong, Russell LoBrutto, Wim F. J. Vermaas

**Author notes:** To whom correspondence should be addressed: School of Life Sciences, Arizona State University, PO Box 874501, Tempe, 85287-4501.

## Abstract

A hypothetical protein encoded by *Synechocystis* sp. PCC 6803 open reading frame *slr0201*shows high sequence similarity to the C subunit of a group of unusual succinate dehydrogenases found in some archaeal species. Slr0201 was originally annotated as HdrB, the B subunit of heterodisulfide reductase, but appears to be SdhC instead. This protein was overexpressed in *E. coli* by cloning the PCR-derived *slr0201* open reading frame into a pET16b-based expression vector. The overproduced Slr0201 accumulated predominantly in inclusion bodies with an apparent molecular mass of 33 kDa. The protein contained at least one [2Fe-2S] cluster based on UV-visible absorbance and CD spectra and EPR spectroscopy, in conjunction with stoichiometric analysis of protein-bound iron and sulfur content. Redox titration showed a midpoint potential (*E*_*m*_) of + 17 mV at pH 7.0, which is consistent with Slr0201 serving a role in transferring electrons between succinate and plastoquinone. Slr0201 was also overproduced in *Synechocystis* sp. PCC 6803 by introducing an additional, His-tagged *slr0201* into the *Synechocystis* genome replacing *psb*A3, creating the *slr0201*^+^-His overexpression strain. Immunoblot analysis shows that Slr0201 is membrane-associated in the wild type. However, in the Slr0201^+^-His strain, immunoreaction occurred in both the membrane and soluble fractions, possibly as a consequence of processing near the N-terminus. The results obtained with Slr0201 are discussed in the light of one of the cyanobacterial SdhB subunits, which shares redox commonalities with archaeal SdhB.

## Introduction

In cyanobacteria, photosynthesis and respiration share several components including electron transfer through the plastoquinone (PQ) pool in thylakoid membranes. Three enzymes, succinate dehydrogenase (SDH), a NADPH-preferring type I dehydrogenase (NDH-1) and a NADH-oxidizing type II dehydrogenase (NDH-2) are potential donors of respiratory electrons into the PQ pool in the model cyanobacterium, *Synechocystis* sp. PCC 6803 (1-3), consistent with open reading frames in the genome of *Synechocystis* sp. PCC 6803 (4). In the case of SDH, the genome of *Synechocystis* sp. PCC 6803 contains two *sdhB*-like ORFs (*sll1625* and *sll0823*) and one *sdhA*-like ORF (*slr1233*). However, a traditional bacterial *sdhC*-like gene is absent. Deletion of the *sdhB*-like ORFs confirmed functional involvement of succinate dehydrogenase in redox poising of the PQ pool in thylakoids (5). Deletion of NDH-1, NDH-2 or SDH genes resulted in a more oxidized PQ pool in darkness, suggesting that these enzymes participate in electron flux into the PQ pool (6). However, SDH appears to catalyze a larger flux of respiratory electrons into the PQ pool than NDH-1 and NDH-2. Moreover, the role of NDH-1 in redox poising of the PQ pool was found to be succinate-mediated, and the effects of NDH-1 deletion were actually attributed to the lack of oxidized NADP required for conversion of isocitrate into 2-oxoglutarate by isocitrate dehydrogenase (6).

Whereas these results confirm the functional presence of SDH in *Synechocystis* sp. PCC 6803 and highlight its contribution to redox poising of the PQ pool in thylakoids, characterization of SDH in *Synechocystis* sp. PCC 6803 is still very incomplete. As indicated, sequence homologues to the membrane anchor subunits (SdhC and/or SdhD) of a typical SDH are missing in the *Synechocystis* genome. However, a hypothetical protein encoded by *slr0201* shows a high sequence similarity to the C subunit of archaeal SDHs. These archaeal SdhC subunits appear to be fused dimers, and have about 10 highly conserved Cys residues (7-9) that are also conserved in Slr0201. In *Sulfolobus tokodaii*, SdhC was found to carry a [2Fe-2S] center (10). Slr0201 was originally annotated as HdrB because of limited sequence similarity to the B subunit of heterodisulfide reductase in some bacteria and archaea. However, we have confirmed Slr0201 to be SdhC based on gene deletion studies (unpublished observations).

To elucidate the redox properties of SdhC and thereby to characterize the role of SdhC in respiratory electron transfer in cyanobacterial thylakoids, in this study we cloned *slr0201* from *Synechocystis* sp. PCC 6803 and overexpressed it in *E. coli*. Biochemical and spectroscopic analysis show that *slr0201* can be overexpressed in *E. coli* and encodes a [2Fe-2S]-containing protein. Moreover, an additional, His-tagged *slr0201* was introduced into *Synechocystis* genome by replacing *psbA3*, one of the two active *psb*A genes coding for the D1 protein of photosystem II (11). In *Synechocystis* sp. PCC 6803, the His-tagged *slr0201* was also overexpressed and the produced full-length Slr0201-His was primarily associated with the membrane fraction.

### Experimental procedures

#### Strains and growth conditions

The *E. coli* strains DH 5α and BL21 (DE3) *plys*S (Stratagene) were used for plasmid propagation and overexpression of the recombinant protein, respectively. Both strains were grown at 37°C in Luria-Bertani (LB) medium containing 50 μg/ml ampicillin or 50 μg/ml chloramphenicol. *Synechocystis* sp. PCC 6803 was grown photomixotrophically at 30°C at a light intensity of 50 μmol photons m^-2^ s^-1^ in BG-11 medium (12) supplemented with 5 mM glucose.

#### *Construction of expression plasmids and transformation of* Synechocystis *sp. PCC 6803*

The open reading frame *slr0201* was amplified by PCR with *Synechocystis* sp. PCC 6803 genomic DNA as a template using the primer 5’-TCCCGTCGCC<u>CAT</u>*<u>ATG</u>*ATTACT GCC-3’ and the reverse primer 5’-GTT<u>GGATCC</u>G*TTA*AGTAAGACCAA-3’, introducing an *Nde* I site (underlined) at the start codon (italicized) and a *Bam*H I site (underlined) adjacent to the stop codon (italicized). Following PCR amplification, the resulting DNA fragment was purified, digested with *Nde* I and *Bam*H I, and separated on a low-melting point agarose gel. The 906-bp DNA fragment was collected and ligated into the plasmid pET16b (Novagen, Madison, WI, USA). The ligation mixture was transformed into *E. coli* strain DH5α, and clones carrying the desired pETS0201 plasmid were identified by restriction digestion analysis. The *slr0201* sequence in the selected clones was verified by DNA sequencing of the entire coding region.

Overexpression of *slr0201* in *Synechocystis* sp. PCC 6803 was accomplished by inserting an additional *slr0201* copy replacing the coding region of *psb*A3, one of two active *psb*A copies. To construct the *Synechocystis* overexpression plasmid, we used the primer pA3S0201a (5’-GTCGCCG<u>TC</u>*<u>ATG</u>*<u>A</u>TTACTGCCCTTGAATAT G-3’) and pA3S0201b (5’-GCCTGAACTAATTGTTA<u>CTGCAG</u>*TTA***GTGATGGTGATGGTGATG**AGTAAGACCAAGC-3’) to amplify the *slr0201* coding region from *Synechocystis* sp. PCC 6803. Introduced restriction sites (*Bsp*H I and *Pst* I) at the start codon and adjacent to the stop codon have been underlined, the start and stop codons have been italicized, and the (His)_6_ tag at the end of the gene has been bolded. The plasmid pA3lhcgA3 (13) was used as overexpression vector that was modified by replacing the gentamycin-resistance cartridge by a spectinomycin-resistance cartridge. The PCR-derived *slr0201*-His (a 924-bp *Bsp*H I-*Pst* I fragment) was cloned into the modified pA3lhcgA3 by restriction digestion removing the modified *lhcb* gene in pA3lhcgA3 at its *Nco* I and *Pst* I sites and ligating with the *slr0201*-His, yielding the recombinant plasmid pA3S0201His. This plasmid carries the up- and downstream regions of *psbA3*, with *slr0201*-His, a T_1_T_2_ transcription terminator sequence, and a spectinomycin-resistance cassette replacing the *psbA3* coding region. The recombinant plasmid was sequenced to confirm the sequence accuracy of *slr0201* and the presence of the 6 x His-tag prior to transformation of *Synechocystis*, which was carried out according to Vermaas *et al*. (14). Putative transformants carrying the additional *slr0201* gene were selected on plates for resistance to 0.5 μg ml^-1^of spectinomycin. Transformants were then subcultured in the presence of increasing concentrations of spectinomycin to allow segregation of the wild-type and mutant genome copies. Segregation was confirmed by PCR using the primers 5’-CATATACATAACCGGCTCCC-3’ and 5’-CTTCGCCACCGCTTC-3’, which are specific for the upstream flanking region of *psbA3* and for *T*_*1*_*T*_*2*_, respectively.

#### *Overexpression of* slr0201 *in* E. coli *and subcellular fractionation*

The expression plasmid pETS0201 with a T7 polymerase-specific promoter was transformed into *E. coli* strain BL21 (DE3) *plys*S cells (Novagen, Madison, WI, USA) that express T7 polymerase upon induction with isopropyl β-D-thiogalactoside (IPTG). Unless specified otherwise, the expression cultures were grown at 37°C in 1 liter of LB broth with shaking until the optical density at 600 nm reached 0.6-0.7. Expression of *slr0201* was induced by addition of IPTG to a final concentration of 1 mM. After a 4-5 h induction, the cell cultures were harvested by centrifugation at 3000 x *g* for 10 min.

The cell pellets were resuspended in 10 ml of lysis buffer [50 mM Tris-HCl (pH 7.5), 150 mM NaCl, 1 mM MgCl_2_, 5 mM DTT and 1 mM freshly prepared PMSF] and lysed by five passages through a French Press cell at 6000 p.s.i. After collecting insoluble components by centrifugation at 30,000 x *g* for 15 min, the supernatant was subjected to ultracentrifugation at 150,000 x *g* for 45 min to sediment membrane parts. The 30,000 x *g* pellet contained inclusion bodies and was washed 3-5 times in washing buffer containing 50 mM Tris-HCl (pH 7.5), 5 mM EDTA, 2 M urea, 2% Triton X-100, and 10 mM DTT. Where indicated, the 30,000 x *g* pellet was then solubilized at room temperature in 50 mM Tris-HCl (7.5) buffer containing 8 M urea, 0.1 M β-mercaptoethanol, 1 mM EDTA, and 1 mM freshly prepared PMSF for 2-3 h with gentle stirring. Cell debris was removed from the solubilized inclusion bodies by centrifugation at 30,000 x *g* for 20 min at 4°C.

#### *Preparation of Slr0201 from* Synechocystis *sp. PCC 6803*

Cell cultures of *Synechocystis* sp.PCC 6803 (1000 ml) with an optical density of 0.6-0.9 at 730 nm were collected and washed. The resulting cell pellets were resuspended in 6 ml of thylakoid buffer containing 20 mM MES-NaOH (pH 6.4), 5 mM MgCl_2_, 5 mM CaCl_2_, 20% glycerol (v/v), 1 mM freshly prepared PMSF, and 5 mM benzamidine. Cell suspensions were transferred into four ice-chilled microcentrifuge tubes filled with 1/3 vol of glass beads prewetted by thylakoid buffer. The cells were broken in a Mini Bead Beater (3110BX, BioSpec Products, Bartlesville, OK) by 6 cycles of shaking; each cycle was 30 s followed by cooling for 3-5 min on ice water. After centrifugation at 1,600 x *g* for 10 min to remove unbroken cells and cell debris, the supernatant was centrifuged again at 45,000 x *g* for 30 min at 4°C to sediment the membrane fraction. The resulting supernatant and the membrane pellet were used as the soluble and membrane protein fraction, respectively.

#### Determination of protein, protein-bound iron, and acid-labile sulfur

Protein content was routinely estimated using bicinchoninic acid-protein assay reagents (Pierce) according to the manufacturer’s instruction with bovine serum albumin as a standard. For stoichiometric analysis of iron-sulfur clusters, the protein content was also determined by the modified Lowry’s method (15). The protein-bound iron and acid-labile sulfur contents in the overexpressed protein samples were determined chemically following the method described by Fish (16) and Rabinowitz (17), respectively. In addition, the protein-bound iron content was determined by atomic absorption spectrophotometry using a Varian SpectrAA 400 Flame Atomic Absorption Spectrometer.

#### Spectroscopic methods

Absorption spectra were recorded with a UV-Vis recording spectrophotometer (UV-2501 PC, Shimadzu). Circular dichroism spectra were recorded on a Jasco 715 CD spectrometer, using a slit width of 2 nm. For EPR spectroscopy measurements, both the 8-M urea solubilized Slr0201 inclusion bodies and the solid Slr0201 inclusion body pellet [directly put into EPR tubes and immersed with 0.3 ml PBS buffer containing 20 mM phosphate (pH 7.4) and 10 mM NaCl] were used. X-band EPR spectra were recorded on a Bruker E580 spectrometer, equipped with an ESR 900 continuous flow liquid helium cryostat (Oxford Instruments). Spectrometer settings were: microwave power: 2 mW (unless indicated otherwise); magnetic field modulation amplitude: 0.3 mT; microwave frequency: 9.41 GHz; magnetic field modulation frequency: 100 kHz. For power saturation studies, the EPR signal at *g* = 1.93 (the *g*_y_ band of [2Fe-2S] clusters) of solid Slr0201 inclusion bodies was recorded at microwave powers varying from 0.001-36 mW at three different temperatures (5, 15 and 25 K). The power for half saturation (*P*_*1/2*_) at each temperature was determined by a fit to a plot of *S* versus log *P* using the equation 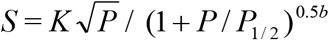, where *S* is the signal amplitude, *K* is a proportionality factor, *P* is the microwave power, *P*_*1/2*_ is the microwave power for half saturation, and *b* is the inhomogeneity parameter (18).

#### Redox titration

The redox midpoint potential of Slr0201 was determined by optical absorbance monitored redox titrations. The Slr0201 solution (∼ 25 μM) in 50 mM Hepes-NaOH (pH 7.0) and 400 mM NaCl was kept anaerobic by flushing the solution with nitrogen gas during the titration. The redox potential of the solution was adjusted by adding freshly prepared sodium dithionite or potassium ferricyanide using a gas-tight 10-μl Hamilton microsyringe. The potentials were measured with a MI-800/4154 microelectrode (Microelectrodes Inc., Bedford, NH) and are quoted relative to the normal standard hydrogen electrode. The microelectrode was calibrated with saturated quinhydrone at pH 4.0 and 7.0, respectively. To facilitate redox equilibration, the following redox mediators were added at a final concentration of 10 μM: safranine 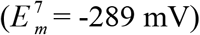, menadione 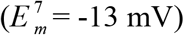, duroquinone 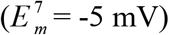, methylene blue 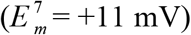 and phenazine methosulfate 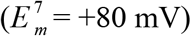.

#### SDS/PAGE, ligand affinity chromatography, and western blotting

For analysis of overexpressed Slr0201 from *E. coli*, the purified, solubilized inclusion bodies were used directly for SDS-PAGE analysis. For analysis of Slr0201 from *Synechocystis* sp. PCC 6803, thylakoid membranes were solubilized in freshly-made sample buffer containing 125 mM Tris-HCl (pH 6.8), 2% SDS, 3% β-mercaptoethanol and 10% (v/v) glycerol at room temperature for about 1 hour. Samples of thylakoids membranes equivalent to 10 μg (for dye staining) or 1 μg (for immunoblotting) of chlorophyll, or of the supernatant (∼3 μg protein for dye staining, and ∼1 μg for immunoblotting) were loaded in each lane and separated on a 10-18% continuous gradient polyacrylamide gel containing 7 M urea. After electrophoresis, proteins on the gel were transferred to a nitrocellulose membrane (pore size 0.45 μm; Schleicher & Schuell, Keene, NH) using the TransBlot Electrophoretic Transfer Cell system (Bio-Rad). Antibody crossreactions were performed essentially as previously described (13). Polyclonal antibodies against Slr0201 were raised in rabbits at Rockland Immunochemicals, Inc. (Gilbertsville, PA) using as antigen the purified, solubilized Slr0201 inclusion bodies that were produced by the *E. coli* strain BL21 (DE3) *plys*S carrying the pETS0201 plasmid. The anti-Slr0201 antiserum was purified by affinity chromatography on an affigel-5 (Biogel) column, to which Slr0201was coupled according to the manufacturer’s instructions. Goat anti-rabbit IgG (H + L) alkaline phosphatase conjugates were used as a secondary antibody (BIO-RAD, 1:3,000). For antibody reaction again the His-tag, we used a commercial monoclonal His-tag antibody (Novagen, 1:2,000), and goat anti-mouse IgG (H + L) alkaline phosphatase conjugates were used as a secondary antibody (BIO-RAD, 1:2,500). Final color development was carried out in alkaline phosphatase buffer [25 mM Tris-HCl (pH 9.8), 150 mM NaCl, and 5 mM MgCl_2_] containing the chromogenic substrates 5-bromo-4-chloro-3-indolyl phosphate (0.015%) and nitroblue tetrazolium (0.03%).

## Results

### *Sequence alignment of* slr0201

The deduced amino acid sequence of Slr0201 shows a high similarity with the C subunit of a group of novel succinate dehydrogenases (SDH) in some archaea. Homologues are also found in several cyanobacteria and in the bacterium *Campylobacter jejunii* (Fig. 1). No homologue of a typical bacterial or mitochondrial SDH membrane anchor is apparent in the *Synechocystis* genome. Slr0201 has ten highly conserved cysteine residues, arranged in two tandem repeat units that each spans half of the protein. Such cysteine-rich motifs may be associated with cofactors such as iron-sulfur clusters.

**Figure 1.**
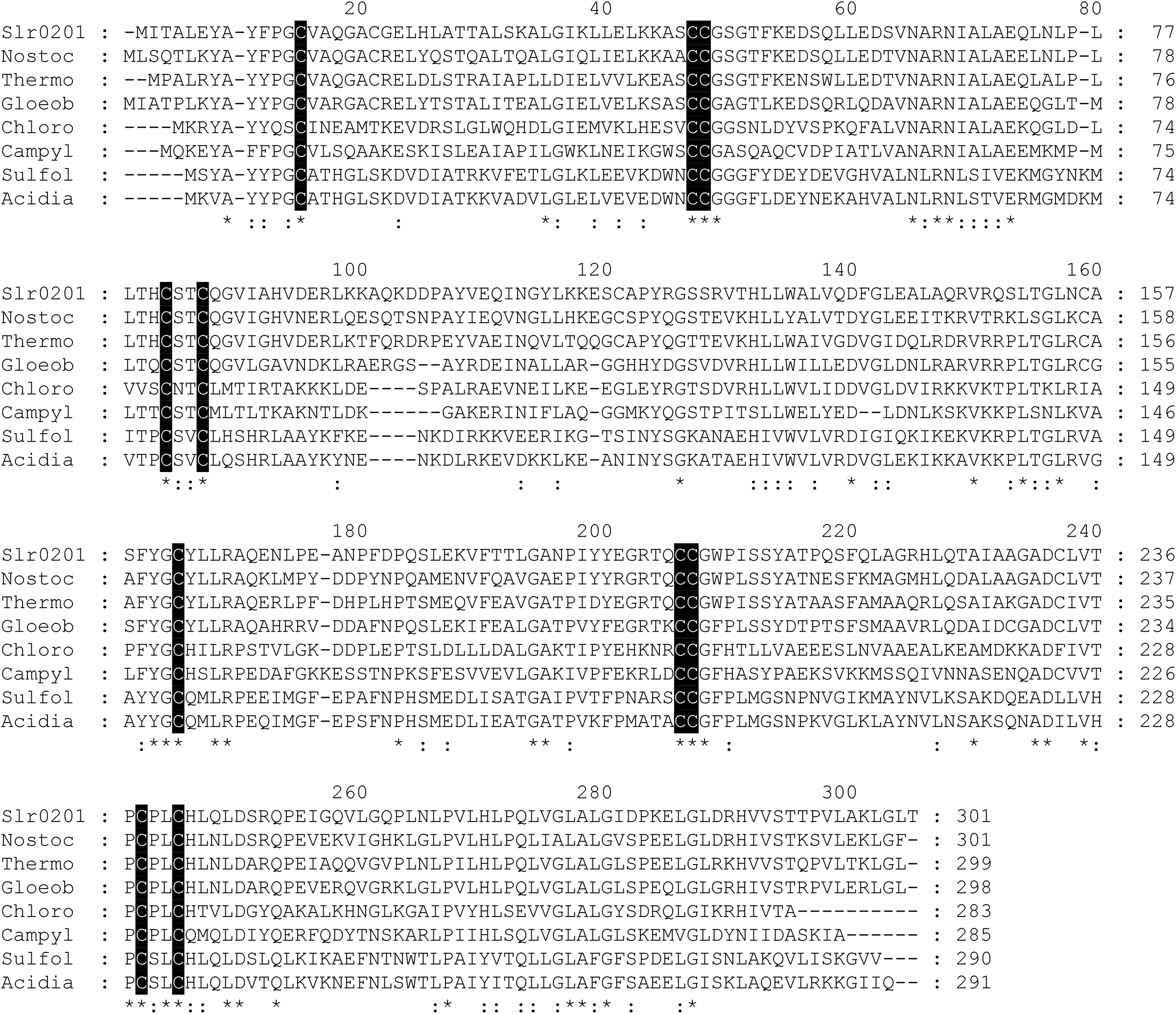
Multiple amino acid sequence alignment of Slr0201 with homologues from *Nostoc* sp. PCC 7120 (Nostoc), *Thermosynechococcus elongatus* BP-1 (Thermo), *Gloeobacter violaceus* PCC 7421 (Gloeob), *Chlorobium tepidum* (Chloro), and *Campylobacter jejuni* (Campyl), and SdhC from *Sulfolobus acidocaldarius* (Sulfol) and *Acidianus ambivalens* (Acidia). Conserved Cys residues have been shaded; identical and similar amino acid residues in the alignment are indicated by asterisks (*) and colons (:), respectively.

### *Overexpression of Slr0201 in* E. coli

For heterologous overexpression of the full-length, *slr0201* encoding protein in the *E. coli* BL21 (DE3)-*plys*S strain, a pET16b-based overexpression plasmid was constructed (see Materials and Methods). As depicted in Fig. 2, Slr0201 was induced by IPTG. The overproduced Slr0201 has an apparent molecular mass of 35 kDa (including a 9 x His Tag and another ∼ 11 amino acids motif from the pET16b, accounting for an additional ∼ 2kDa) as estimated by SDS-PAGE, which agrees well with the 33.4 kDa calculated molecular weight deduced from the amino acid sequence.

**Figure 2.**
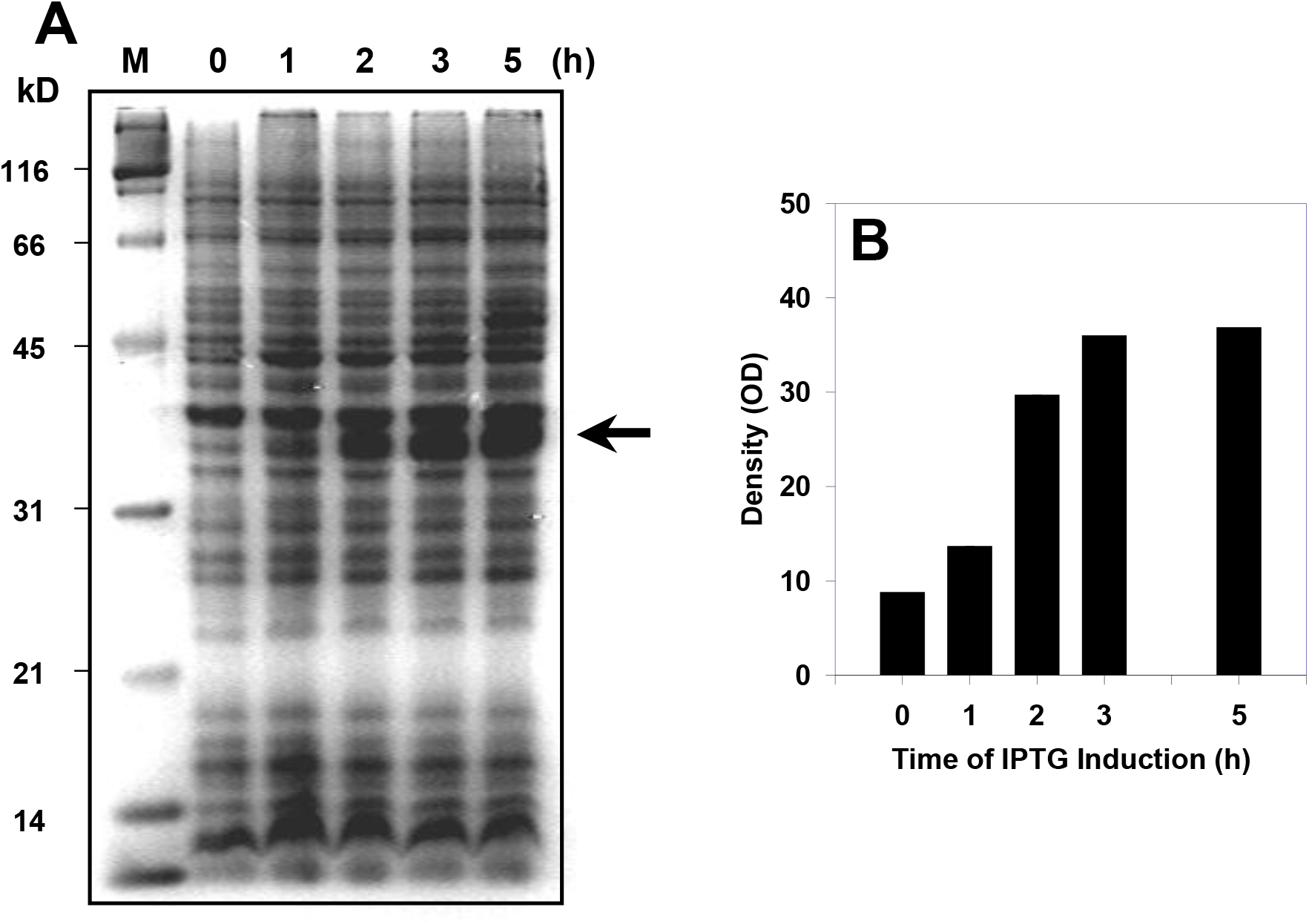
Heterologous overexpression of *slr0201* carrying a 9 x His-tag in *E. coli* strain BL21 (DE3) *plysS* under the T7 RNA polymerase promoter. Overexpression of *slr0201* was induced by addition of 1 mM IPTG at time 0. **A**: Protein profile of the whole cell lysate extracted at the times indicated after IPTG addition. The overexpressed Slr0201 has been indicated by an arrow. M: molecular size marker. **B**: Staining intensity of the Slr0201 region as a function of time after IPTG addition.

Upon fractionating the whole cell lysate, the overexpressed s*lr0201* product was found primarily in the fraction containing the inclusion bodies although a minor part of Slr0201 was found in the membrane fraction (data not shown; however, see Fig. 3). The inclusion bodies were easily purified by 3-5 washes to remove membrane lipids, soluble proteins, nucleic acids and cell debris. One liter of cell culture could yield *ca*. 300-500 mg (wet weight) of purified inclusion body pellet, which was solubilized efficiently by incubation with 8 M urea. Slr0201 from the solubilized inclusion bodies that are enriched in the overexpressed protein (> 85%) was purified and extracted from the SDS-PAGE gel, and was used to raise Slr0201 antibodies.

**Figure 3.**
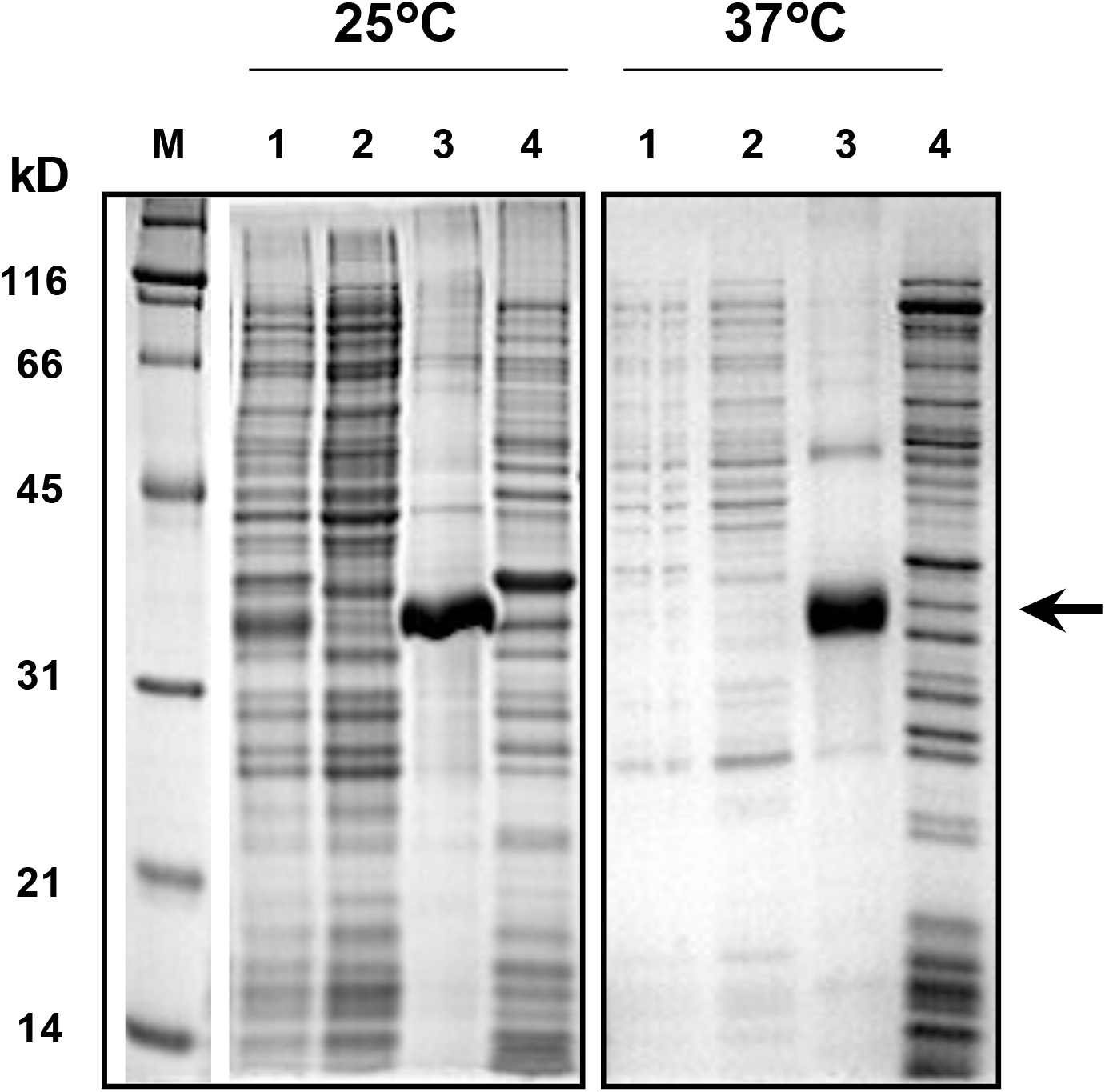
Distribution of Slr0201upon overexpression of *slr0201* in *E. coli* at 25°C and 37°C. The SDS-PAGE gel stained with Coomassie Brilliant Blue shows polypeptide profiles of the whole cell lysate (1), the supernatant (2), the inclusion body pellet (3), and the membrane fraction (4). Slr0201 in the inclusion body fraction (lane 3 in each panel) has been indicated by arrows. Molecular weights of the markers (M) have been indicated.

### *Effects of growth conditions on overexpression of Slr0201 in* E. coli

The fact that overexpressed Slr0201 was found primarily in the inclusion body fraction probably signifies the lack of proper folding of Slr0201 to a native form (19, 20). To investigate whether modification of growth conditions (*e*.*g*., addition of Fe, S and cysteine, as Slr0201 may bind iron-sulfur center) may affect the yield and/or solubility of the overexpressed protein, the transformed *E. coli* strain was grown in the presence of supplementary Fe^+3^ (ferric ammonium citrate, 1 mM), sulfur (MgSO_4_, 1 mM) and cysteine (0.25 mM), and at lower temperature (∼ 25°C). As shown in Fig. 3, the yield of expressed Slr0201 was somewhat improved by lowering the growth temperature, but the expressed protein remained primarily in inclusion bodies. Moreover, providing extra Fe^+3^, sulfur and cysteine did not alter the distribution or abundance of Slr0201 (data not shown).

### Spectral properties of the overexpressed Slr0201

The purified, solubilized inclusion bodies initially showed a dark-brown color suggesting the presence of a bound cofactor. The absorption spectrum (Fig. 4A) was characteristic of [2Fe-2S] cluster-containing proteins that have an absorption peak at 420 nm with a lower, broad shoulder around 550 nm. Such spectral characteristics were fully bleached upon reduction by dithionite. The near-UV/visible CD spectrum of the overexpressed Slr0201 protein showed two negative troughs at 337 and ∼605 nm, a dominant positive peak at 460 nm, and minor positive peaks at 316, 360 and around 620 nm (Fig. 4B). These features are also very similar to [2Fe-2S] clusters from plants (21) although the positions of these dominant positive and negative features are slightly blue-shifted in the overexpressed Slr0201.

**Figure 4.**
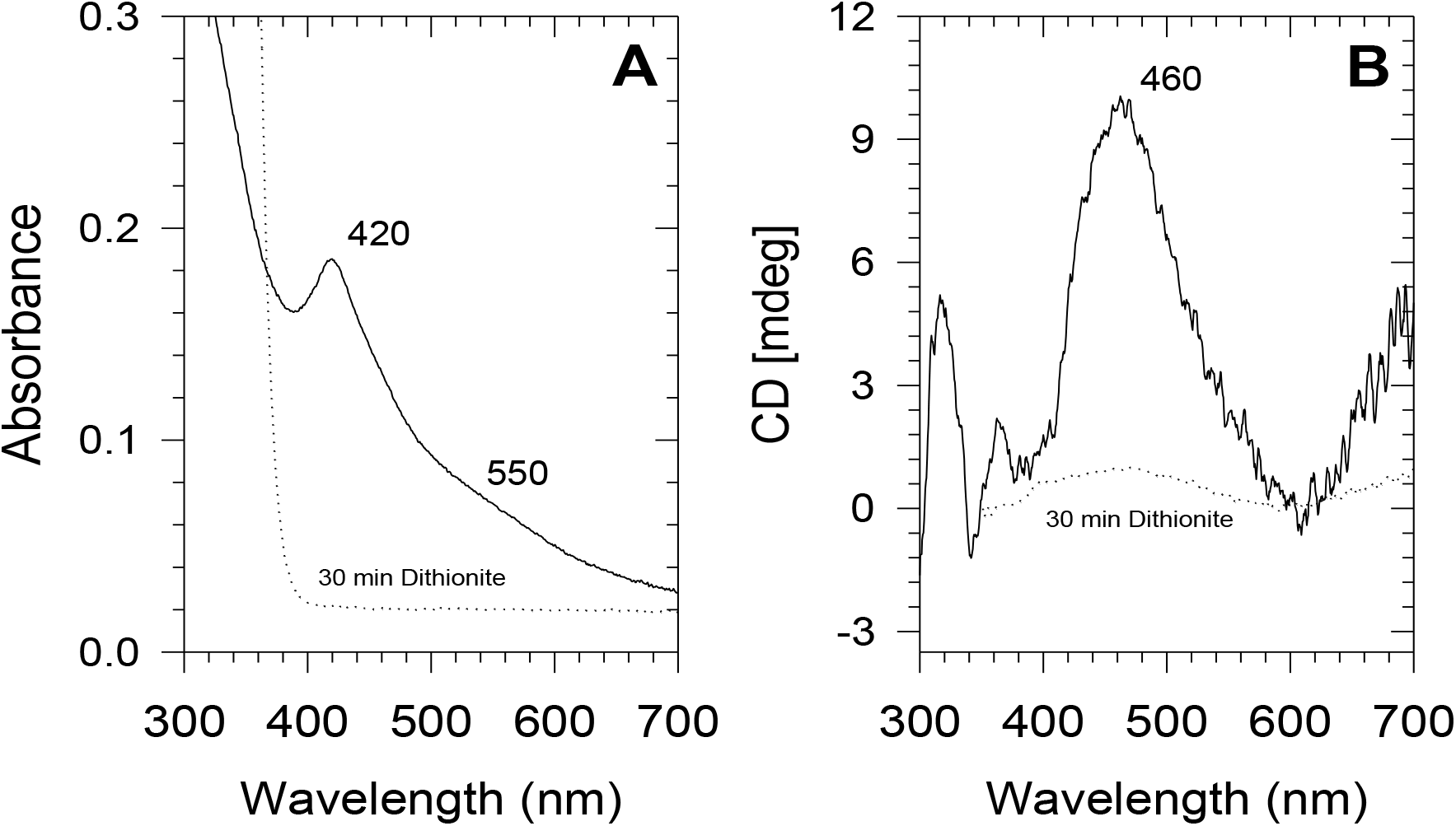
Absorbance (**A**) and CD spectra (**B**) of the Slr0201 inclusion bodies solubilized in Tris-HCl (pH 8.2) containing 8 M urea before (solid line) and 30 min after addition of dithionite (dashed line). The spectra were recorded at 25°C in solubilization buffer at a protein concentration of ∼ 45 μg ml^-1^.

### Redox titration

The potential presence of [2Fe-2S] clusters in Slr0201 was further examined by determination of the midpoint potential via a redox titration monitoring absorbance at 400 nm. A typical redox titration curve of Slr0201 at pH 7.0 is shown in Fig. 5. The data points were fitted with a single Nernstian curve with n = 1. The midpoint potential (*E*_*m*_) of the [2Fe-2S] cluster in Slr0201 under these conditions was estimated to be + 17 mV.

**Figure 5.**
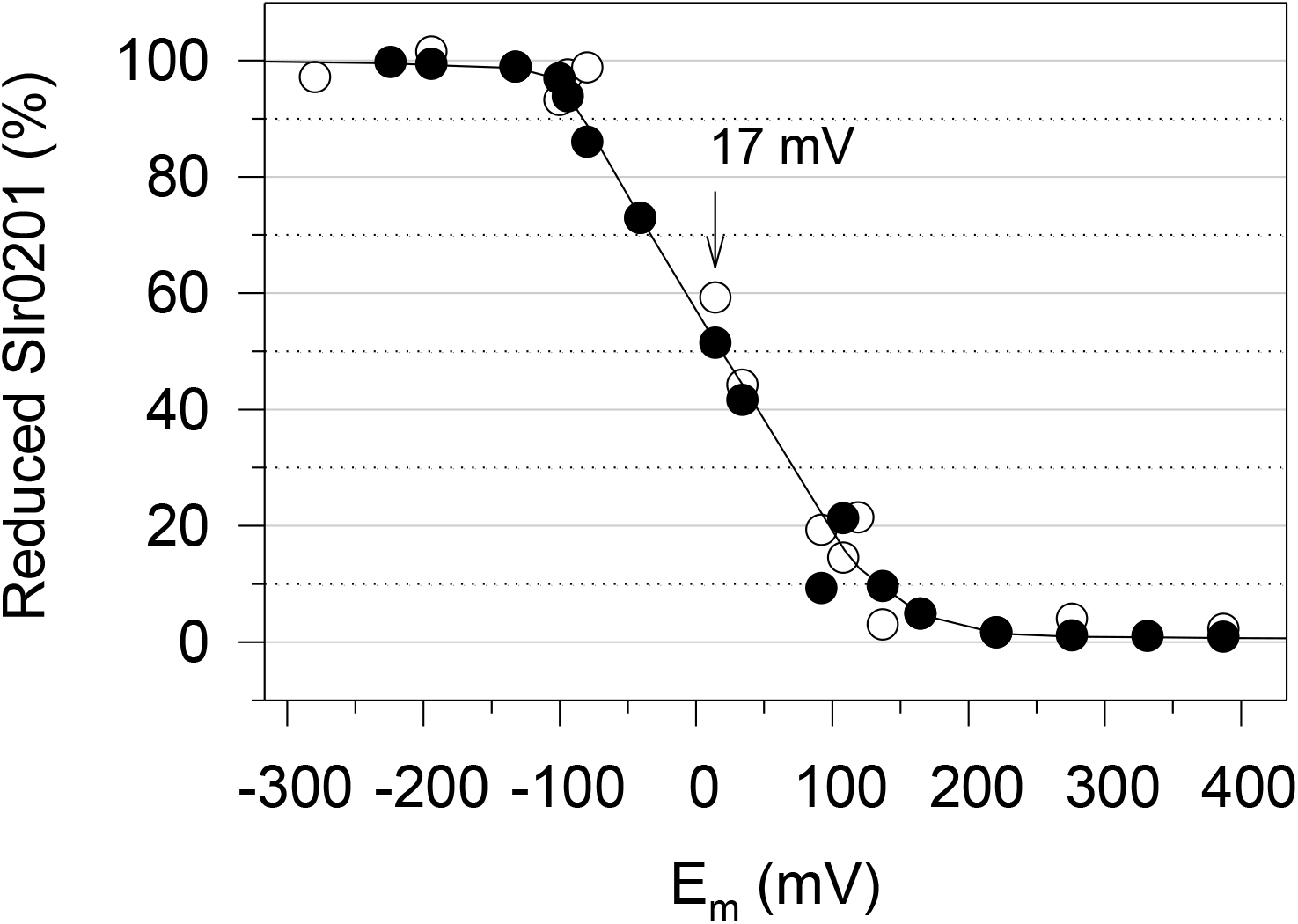
Determination of the midpoint potential of solubilized Slr0201 inclusion bodies by monitoring absorbance at 400 nm as a function of the redox potential. Redox titration was conducted under anaerobic condition in the presence of redox mediators (see Material and Methods). Results are from two separate redox titrations (○ and ●). The data were fit to the Nernst equation for a one-electron couple. The arrow indicates the calculated midpoint potential.

### Chemical analysis of the protein-bound iron and sulfur content in Slr0201

Stoichiometric analysis of protein-bound iron and acid-labile sulfur content showed that the purified inclusion bodies contained an average 2.03 ± 0.3 iron atoms, and 1.95 ± 0.2 sulfur atoms per monomer of Slr0201.

### EPR spectroscopy

The 8-M urea-solubilized, air-oxidized Slr0201 inclusion bodies did not elicit iron-sulfur specific EPR signals with typical *g* values in range of 1.8 ∼ 2.1. An EPR signal at *g* = 4.3 was observed, but this was attributed to free, high-spin Fe^+3^ in a rhombic environment (Fig. 6A). While a similar EPR spectrum was observed with the dithionite-reduced, solubilized Slr0201, a weak EPR signal at g ∼1.93 was detected (Fig. 6B). However, using the solid Slr0201 inclusion body pellet directly without urea solubilization, a [2Fe-2S]-typical rhombic resonance signal with values of *g*_z_ = 2.03, *g*_med_ = 1.934, and *g*_min_ = 1.907 was recorded (Fig. 6C), although reduction by dithionite did not result in further development of this resonance (Fig. 6D).

**Figure 6.**
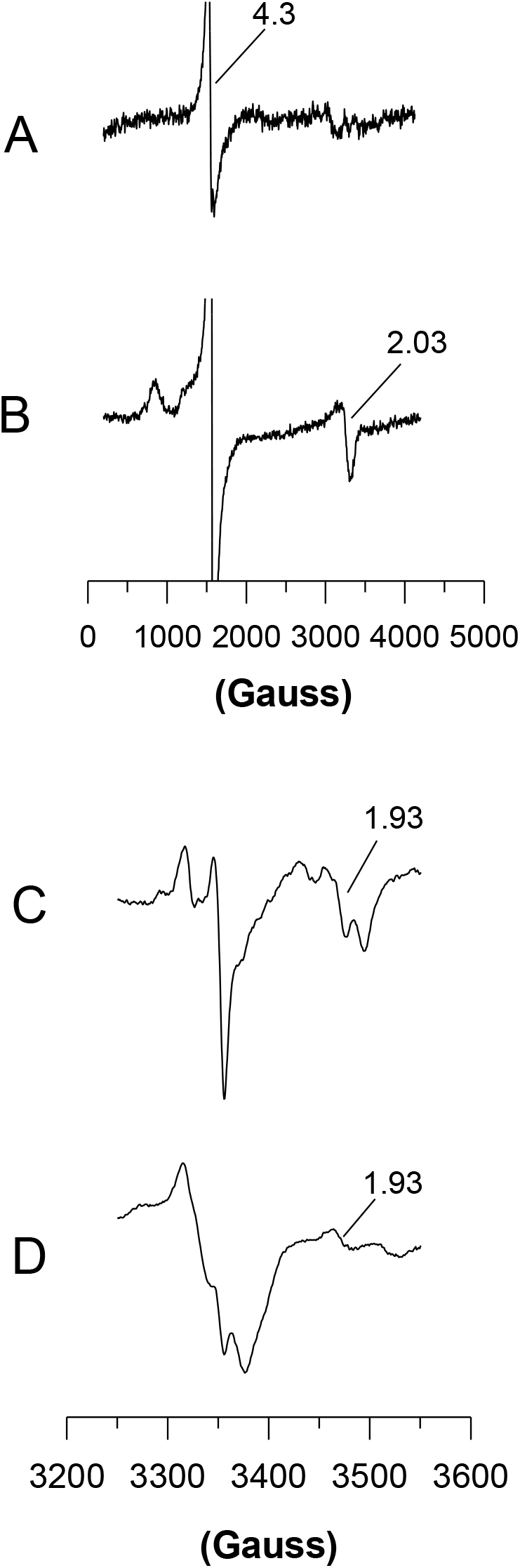
EPR spectra of solubilized Slr0201 inclusion bodies (**A**: as prepared; **B**: reduced with dithionite) and a solid Slr0201 inclusion body pellet (**C**: as prepared; **D**: reduced with dithionite) at 15 K. Other condition settings for EPR measurements include: microwave power: 0.21 mW; microwave frequency: 9.4156 GHz; modulation amplitude: 5 G; gain: 60 dB; time constant: 0.1638 s; sweep time: 41.94 s; and number of scans: 16.

[4Fe-4S] clusters may also exhibit a rhombic resonance signal with *g* values very similar to those from [2Fe-2S] cluster (22, 23). However, compared to those from [2Fe-2S] clusters, the EPR resonance from [4Fe-4S] clusters shows a very different relaxation behavior. EPR resonance signals from [4Fe-4S] clusters become undetectable over 35-50 K, due to substantial broadening effects at higher temperatures (24). The FeS center EPR resonance from Slr0201 was observed optimally at temperatures around 15 K (Fig. 7), and was still detectable at temperatures over 70 K (data not shown), a feature typical of [2Fe-2S] clusters. The power saturation behavior was also well in agreement with those observed in [2Fe-2S] clusters (25): the computed *P*_*1/2*_ values were ∼0.05, 0.35, and 2.2 mW at 5, 15, and 25 K, respectively (Fig. 8).

**Figure 7.**
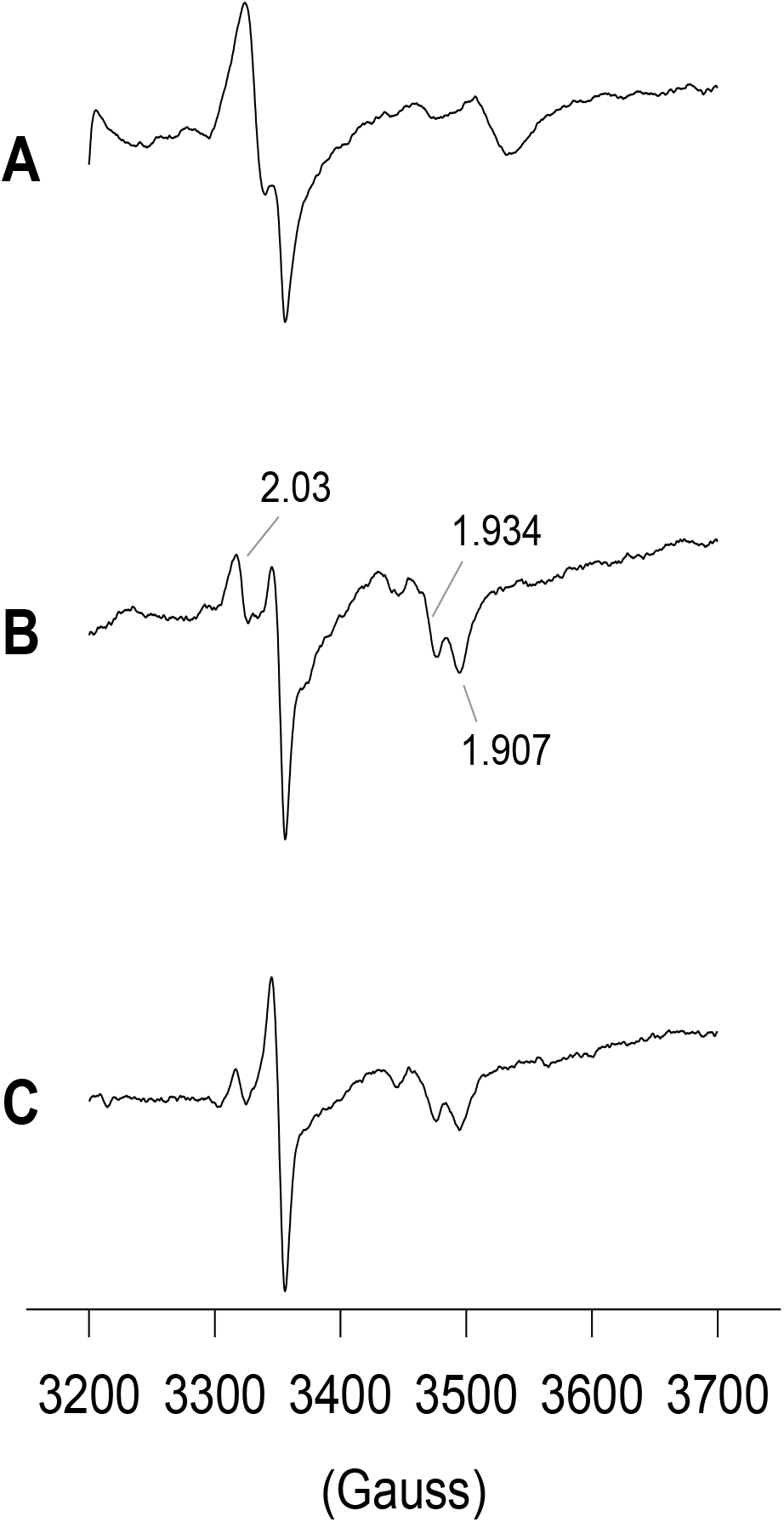
EPR spectra of the solid Slr0201 inclusion body pellet detected at three different temperatures (**A**: 5 K; **B**: 15 K; **C**: 25 K). All other conditions are as in Fig. 6.

**Figure 8.**
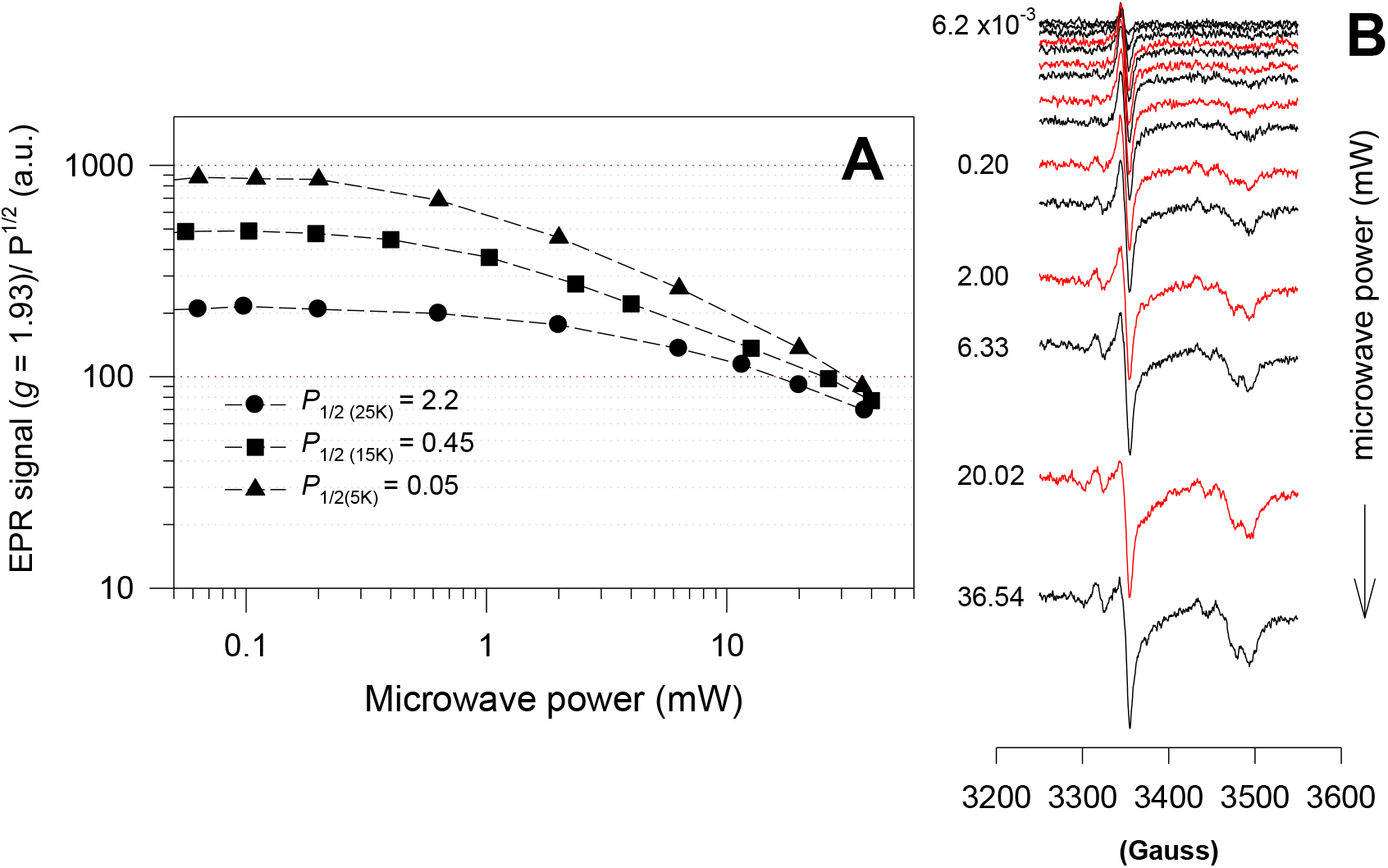
(**A**). Power saturation curves of the *g* = 1.93 EPR signal of the Slr0201 inclusion body pellet detected at 5, 15 and 25 K. (**B**). EPR spectra of the solid Slr0201 inclusion body pellet detected at different microwave power levels at 5 K.

### *Overexpression of Slr0201 in* Synechocystis *sp. PCC 6803*

To investigate whether Slr0201 could be overexpressed in *Synechocystis* sp. PCC 6803, an additional copy of *slr0201* was introduced into the *Synechocystis* genome by replacing *psbA3*, one of the two active *psbA* copies. The resulting *Synechocystis* strain was named *slr0201*^+^-His. As shown in Fig. 9, immunoblotting analysis using antibodies raised against the purified, solubilized Slr0201 inclusion bodies detected one specific protein band with an electrophoretic migration consistent with Slr0201 in both the wild type and the *slr0201*^+^-His strain. The crossreaction occurred exclusively in the membrane fraction (Fig. 9C), which is indicative of an association of Slr0201 with membranes *in vivo* in *Synechocystis* sp. PCC 6803. The immunoreaction intensity of the Slr0201 band detected in the *slr0201*^+^-His strain was significantly higher than that in the wild type, indicating that, as expected, more Slr0201 was produced in the overexpression strain. However, no clear differences are apparent in the Coomassie-stained gel, suggesting that enhanced expression of Slr0201, if any, is moderate in the *slr0201*^+^-His strain. As expected, using the monoclonal His-tag antibody, the specific protein band was detected only in the *slr0201*^+^-His strain (Fig. 9D), indicating that the inserted, additional copy of *slr0201* was expressed in *Synechocystis* sp. PCC 6803. However, the His-tag antibody-mediated crossreaction occurred not only in the membrane fraction but also in the soluble fraction, where a protein band with a slightly lower molecular size (∼31 kDa) was detected.

**Figure 9.**
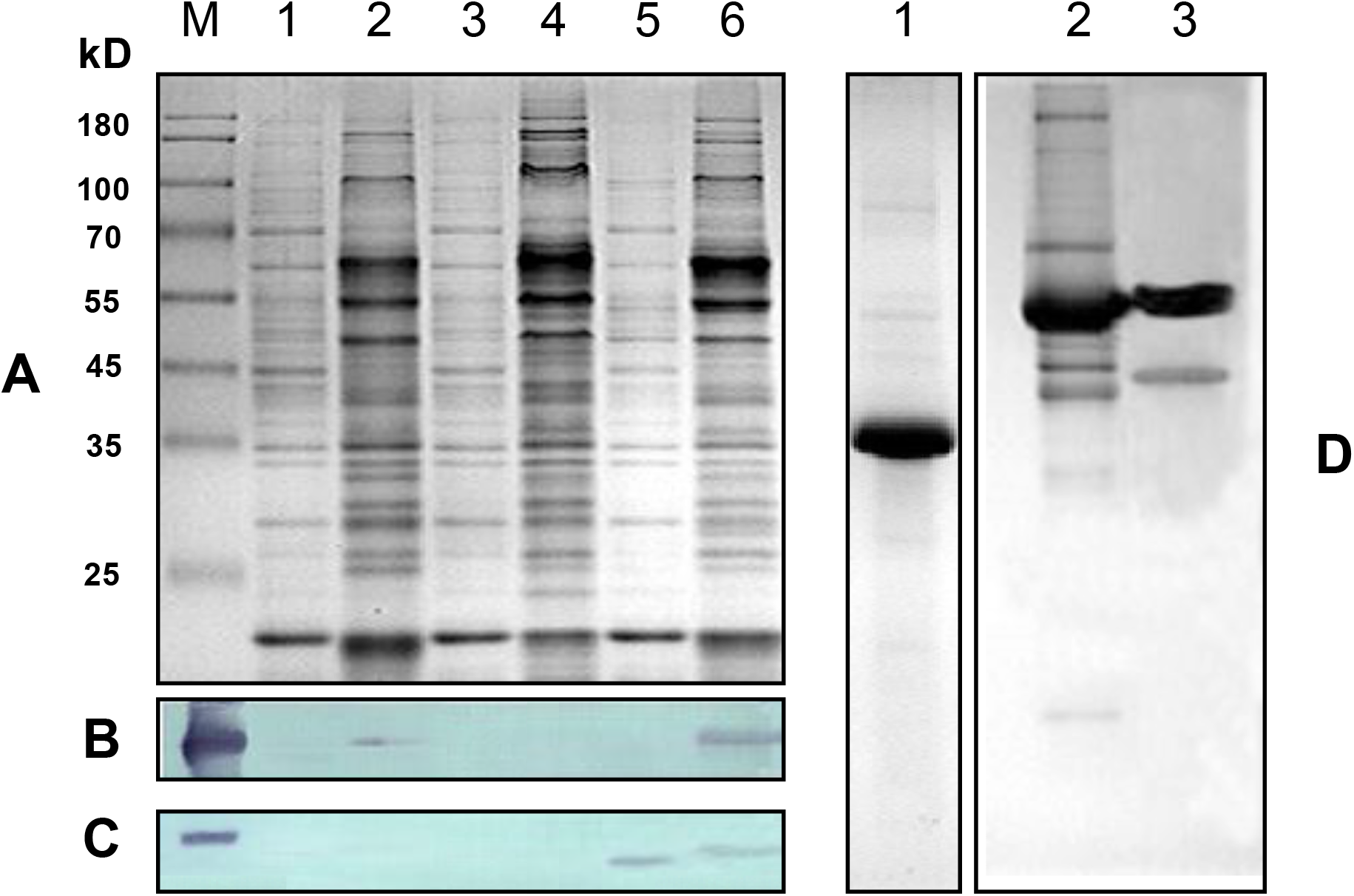
Analysis of expression of *slr0201* in *Synechocystis* sp. PCC 6803. Soluble (1, 3, 5) and membrane proteins (2, 4, 6) from the wild type (1, 2), the Δ*slr0201*mutant (3, 4), and the *slr0201*^+^-His strain (5, 6) were separated by SDS-PAGE gel (**A**). After transfer to nitrocellulose membrane, the expressed *slr0201* products were immunoreacted with either antibodies raised against Slr0201 (**B)** or a commercial His-tag monoclonal antibody for expression of the inserted, His-tagged *slr0201* (**C**). The sample loaded in lane “M” in panel **B** and **C** is the purified Slr0201 overproduced in *E. coli* strain BL21 (DE3) *plysS* carrying the plasmid pET16S0201. Panel **D** shows the purified, solubilized Slr0201 inclusion bodies used as antigen to raise Slr0201 antibodies (lane 1) as well as the Slr0201 antibodies before (lane 2) and after (lane 3) affinity purification.

## Discussion

### *Heterologous overexpression of* slr0201 *in* E. coli

Similar to other *Synechocystis* cofactor-containing proteins overproduced in *E. coli* (*e*.*g*., 26-27), the recombinant Slr0201 overproduced in *E. coli* accumulates predominantly in inclusion bodies. The poor solubility of overproduced Slr0201 does not seem to be due to limiting iron and/or cysteine, and may be ascribed to aggregation and/or misfolding during overexpression. The main protein from the purified, solubilized inclusion bodies migrates on SDS-PAGE at ∼35 kDa (including ∼ 2 kDa from the vector), an apparent molecular weight corresponding closely to the 33.4 kDa weight deduced from the sequence data (Fig. 1) and very similar to that of the C subunit of a group of archaeal SDHs found in *Sulfolobus acidocaldarius* (32.1 kDa, 7), *Acidianus ambivalens* (33 kDa, 9), and *Sulfolobus tokodaii* (33 kDa, 10).

### *The* slr0201 *gene encodes an [2Fe-2S]-cluster containing protein*

Sequence alignments show that Slr0201 contains Cys-rich motifs (Fig. 1), and the conservation of cysteine residues suggests that these residues may be important for the function of Slr0201. One possibility is that these conserved cysteine residues provide disulfide bridges linking adjacent subunits. They also may serve as cofactor ligands, most likely of iron-sulfur clusters. Indeed, several lines of evidence provided in this study suggest that *slr0201* encodes a polypeptide containing [2Fe-2S] clusters. First of all, the Slr0201-enriched inclusion bodies overproduced in *E. coli* show a dark brown color with optical absorbance and CD spectra (Fig. 4) resembling those observed with [2Fe-2S]-ferredoxin-type proteins (*e*.*g*., 21, 28, 29). Secondly, the content of protein-bound iron and acid-labile sulfur (2.03 ± 0.3 iron and 1.95 ± 0.2 sulfur per monomer of the Slr0201) corroborates the spectroscopic results, and supports the presence of iron-sulfur clusters in Slr0201. Moreover, the solid Slr0201 inclusion body pellet provided a rhombic EPR resonance with typical *g* values characteristic of [2Fe-2S] clusters (Figs. 6 and 7). In addition, power saturation and temperature-dependent relaxation behaviors of the observed EPR resonance (Figs. 7 and 8) are all characteristic of EPR signals derived from [2Fe-2S] clusters.

It is noteworthy that sequence alignments with Slr0201 show two tandem repeat units containing cysteine motifs (each contains five highly conserved cysteine residues, Fig. 1), suggesting a gene duplication event (9) and the presence of two iron-sulfur clusters per polypeptide. However, the FeS/protein stoichiometry of the inclusion bodies suggests the presence of only one [2Fe-2S] cluster per polypeptide. In previous iron-sulfur protein studies, the results of chemical analysis sometimes did not match the expected number of protein-bound iron-sulfur clusters. For example, chemical analysis of the iron and sulfur contents in LipA, a [3Fe-4S]- and a [4Fe-4S]-cluster containing enzyme that is involved in lipoic acid biosynthesis, gave 3.4 ± 0.4 iron and 4.8 ± 0.8 sulfur atoms per LipA monomer (30). In an *in vitro* dissociation experiment, Ugulava *et al*. (31) reported a ratio of 2.4 ± 0.5 iron atoms per biotin synthase dimer containing two [2Fe-2S] clusters. Lower iron and/or sulfur contents in iron-sulfur proteins were probably due to (1) apoproteins without iron-sulfur cluster present in the sample (32); and/or (2) loss of the iron-sulfur cofactor during isolation, especially if high concentrations of urea were used (33).

Considering the possibility that the iron-sulfur clusters in Slr0201 may not be completely assembled when the protein is overproduced as inclusion bodies in *E. coli*, and some of the iron-sulfur clusters may have been lost during solubilization of the Slr0201 inclusion bodies with 8 M urea, we suspect that our chemical analysis data may underestimate the iron-sulfur content of Slr0201. A final resolution of the number of [2Fe-2S] clusters in Slr0201 awaits a more detailed examination.

Indeed, [2Fe-2S] clusters in Slr0201-containing inclusion bodies were unstable when incubated with 8 M urea for 4-6 h at room temperature. During this time period, the brown color gradually disappeared and the solubilized inclusion bodies became EPR silent except for a *g* = 4.3 signal (characteristic of high-spin Fe^3+^; Fig. 6A and B). Since the solid inclusion body pellet exhibits an EPR resonance typical of [2Fe-2S] clusters, 8 M urea is interpreted to denature Slr0201, leading to decomposition of the iron-sulfur clusters during incubation. This is consistent with observations by Ohmori *et al*. (34) who showed that iron-sulfur clusters in spinach ferredoxin were decomposed over a 4-h incubation period with 4 M urea. No further development of the [2Fe-2S]-special EPR resonance (*g* = 1.93) was observed (Fig. 6) in the dithionite-treated Slr0201 inclusion bodies, which is not unexpected since reduction of the solid Slr0201 inclusion bodies is much less effective compared to a protein solution.

### *Overexpression of Slr0201 in* Synechocystis *sp. PCC 6803*

To elucidate its role *in vivo*, Slr0201 was also overproduced in *Synechocystis* sp. PCC 6803 by introducing an additional, His-tagged *slr0201* into the genome of *Synechocystis* sp. PCC 6803 under the control of the *psb*A3 promoter. The successful integration of the introduced *slr0201* into the genome of *Synechocystis* sp. PCC 6803 was confirmed by DNA sequencing of the *slr0201*^+^-His strain (data not shown). Moreover, immunoblot analysis showed that the introduced *slr0201* was functionally expressed *in vivo* as crossreaction with His-tag antibodies was exclusively detected in the *slr0201*^+^-His strain (Fig. 9). However, immunoreaction occurred not only in membranes but also with a ∼31 kDa band in the soluble fraction, indicating that at least a fraction of the expressed Slr0201 *in vivo* was not associated with membranes. Hydropathy analysis of Slr0201 indicates that this protein is hydrophilic with two uncharged stretches near its N-terminus (the residues 55-78) and C-terminus (the residues 258-280), respectively. This implies that Slr0201 is most likely peripherally associated with membranes, which is consistent with our observation that Slr0201 was easily extracted during membrane isolation. Since the protein band that was positively detected with His-tag antibodies in the soluble fraction was smaller than the Slr0201 associated with membranes, the Slr0201 detected in the soluble fraction may be cleaved near its N-terminus as observed with cytochrome c_M_ in *Synechocystis* sp. PCC 6803 (35). Apparently, such a cleavage alters the ability to attach to membranes and appears to result in a loss of the Slr0201 antibody recognition site, which would explain that only the His-tag antibody crossreaction occurred in the soluble fraction at a smaller molecular size.

### Evolutionary implications

A typical bacterial SDH contains three iron-sulfur clusters: center 1 ([2Fe-2S]); center 2 ([4Fe-4S]); and center 3 ([3Fe-4S]). These iron-sulfur centers are all located in the SdhB subunit, functioning to mediate electron transfer from the SdhA subunit to the membrane anchoring subunit that usually is devoid of [Fe-S] centers (36, 37). However, in the archaeon *Sulfolobus tokodaii* (10, 38), spectrometric and chemical analysis showed an atypical SdhC that strongly resembles Slr0201 (Fig. 1) and that contained a [2Fe-2S] cluster(s). Interestingly, in the atypical SDHs found in some archaea (7-10), the [3Fe-4S] cluster in SdhB (center 3) is replaced by a second [4Fe-4S] cluster, which is accompanied by incorporation of a fourth Cys residue in the cysteine cluster III of SdhB. In *Synechocystis* sp. PCC 6803, one of the SdhB genes (*sll1625*) contains an additional Cys residue (Cys 214) in the cysteine cluster III. This suggests that in these archaeal-type SDHs, the SdhB replacement of a [3Fe-4S] cluster by an additional [4Fe-4S] cluster is functionally coupled to the presence of [2Fe-2S] cluster(s) in SdhC as it has been proposed that the [3Fe-4S] cluster is directly involved in electron transfer to quinones (37, 38).

While the [2Fe-2S]-containing SdhC from *Sulfolobus tokodaii* strain 7 was unstable upon dithionite reduction and ferricyanide re-oxidation, making this protein unsuitable for redox titration and determination of the midpoint potential (10), Slr0201 was stable during the redox titration and the midpoint potential (*E*_*m*_) for [2Fe-2S] clusters in this protein was determined to be +17 mV (pH 7.0) (Fig. 5). This *E*_*m*_ is close to the midpoint potential (*E*_*m*_ = +10 mV) of center 1 [2Fe-2S] in SdhB from *E. coli* (38, 39), and comparable to that of the [2Fe-2S] cluster from SdhB of archaeal SDH as well (7-9).

The membrane-associated Slr0201 is expected to interact with quinones in the membrane. In cyanobacteria, plastoquinone (PQ-9) is the major electron carrier mediating both photosynthetic and respiratory electron transport (3, 40). The *E*_*m*_ of PQ-9 is + 65-100 mV, which is consistent with a role of Slr0201 in electron transfer from SdhB to plastoquinone. Therefore, Slr0201 in *Synechocystis* sp. PCC 6803 may have a function that is fully homologous to that of membrane-associated subunits of SDH from eubacterial and mitochondrial systems, in spite of a very different evolutionary origin.

## Acknowledgment

We thank Dr. Dan Brune for his assistance in the Slr0201 CD spectral measurements, Dr. James Allen for his advice regarding redox titrations, and Tom Colella for his excellent technical assistance in determinations of the protein-bound iron content using a flame atomic absorption spectrophotometer.

## For Table of Contents Use Only

The *Synechocystis* sp. PCC 6803 open reading frame *slr0201* that is homologous to *sdhC* from Archaea codes for a [2Fe-2S] protein.

Fusheng Xiong, Russell LoBrutto and Wim Vermaas

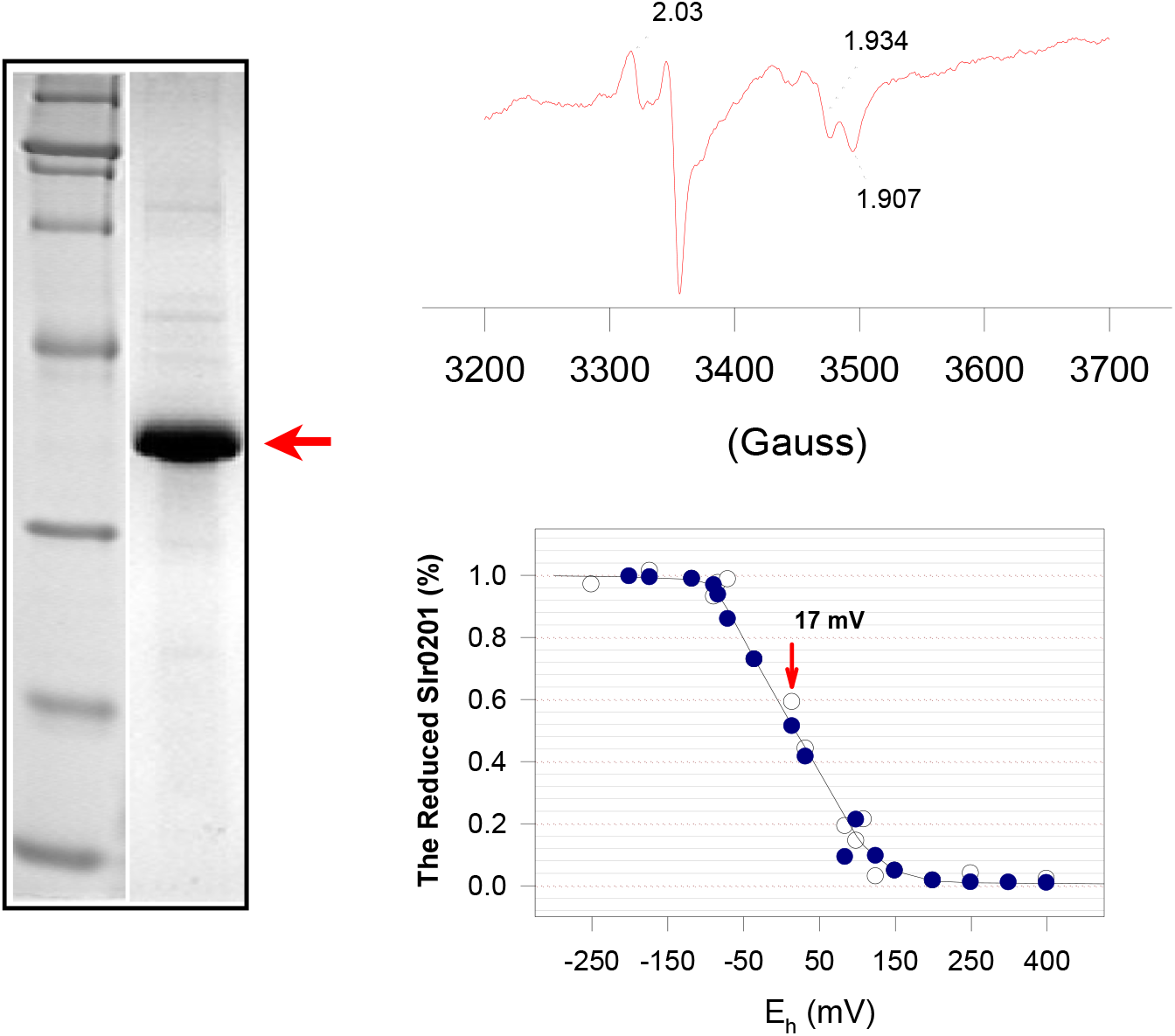

